# Meta-analysis of exhausted CD8+ T cells from *Homo sapiens* and *Mus musculus* provides robust targets for immunotherapy

**DOI:** 10.1101/2021.07.05.451111

**Authors:** Lin Zhang, Yicheng Guo, Hafumi Nishi

## Abstract

T cell exhaustion is a state of T cell dysfunction during chronic infection and cancer. Antibody-targeting immune checkpoint inhibitors to reverse T cell exhaustion is a promising approach for cancer immunotherapy. However, the therapeutic efficacy of known immune checkpoint inhibitors remains low. To expand the potential effective targets to reverse T cell exhaustion, a meta-analysis was performed to integrate seven exhaustion datasets caused by multiple diseases in both humans and mice. In this study, an overlap of 21 upregulated and 37 downregulated genes was identified in human and mouse exhausted CD8+ T cells. These genes were significantly enriched in exhaustion response-related pathways, such as signal transduction, immune system processes, and regulation of cytokine production. Gene expression network analysis revealed that the well-documented exhaustion genes were defined as hub genes in upregulated genes, such as programmed cell death protein 1 and cytotoxic T-lymphocyte associated protein 4. In addition, a weighted gene co-expression analysis identified 175 overlapping genes that were significantly correlated with the exhaustion trait in both humans and mice. This study found that nine genes, including thymocyte selection associated high mobility group box and CD200 receptor 1, were significantly upregulated and highly related to T cell exhaustion. These genes should be additional robust targets for immunotherapy and T-cell dysfunction studies.

## Introduction

T cell exhaustion, a state of differentiation that develops from memory T cells during chronic infections, sepsis, and cancer, is antigen dependent and usually exhibits both poor effector functions in a hierarchical manner and impaired memory T cell potential (1–3). Typically, (i) the robust proliferative and multiple cytokine production capacities that include interferon-γ (IFNγ), tumor necrosis factor (TNF), and interleukin-2 (IL-2) are lost in exhausted cells; (ii) inhibitory checkpoint receptors are expressed in increasing amounts and diversity, such as programmed cell death protein 1 (PD1), lymphocyte activation gene 3 protein (LAG3), hepatitis A virus cellular receptor 2 (HAVCR2/TIM3), cytotoxic T lymphocyte antigen 4 (CTLA4), CD160, 2B4, B- and T-lymphocyte attenuator (BTLA), and T cell immunoreceptor with immunoglobulin and ITIM domains (TIGIT); and (iii) changes in transcriptional profiles and metabolic derangements (1, 2, 4–7). Exhausted T cells maintain their functional exhaustion after exposure to antigens (8). They were initially identified in chronic viral infections in mice(9, 10), and similar dysfunctional features have been observed in cancer(11–13).

Intriguingly, although exhausted T cells are often related to inefficient control of persistent infections and tumors, immune responses may be reinvigorated in exhausted T cells by targeting inhibitory checkpoint receptors (2). For example, the function of exhausted CD8+ T cells present in mice during persistent lymphocytic choriomeningitis virus (LCMV) infection is restored by blocking the interaction of PD1 with its ligand PD-L1 (14). Likewise, blockade of the negative checkpoint receptor CTLA4 allows transient reprogramming and functional rescue of T cell dysfunction due to the tumor microenvironment (11). Of note, antibody blockade of immune checkpoints is considered a promising approach for cancer immunotherapy and therapeutic intervention of virus- or tumor-specific T cells during chronic viral infections or tumors (11, 15, 16). Fortunately, certain antibodies have been developed with good clinical performance; for example, the antibody ipilimumab, targeting the CTLA4 pathway, has achieved US Food and Drug Administration approval as the first immunotherapy for the treatment of melanoma (16). However, some antibodies that block immune checkpoint pathways are still being investigated for their clinical efficacy in patients with melanoma, renal and lung cancers, or multiple cancers, such as the antibodies MDX-1106 targeting PD1 and IMP321 targeting LAG3 (16). However, it is important to emphasize that immunotherapeutics remain problematic. For example, exhausted T cells manifest heterogeneous populations that include T-bet^hi^ PD1^mid^ and eomesodermin (EOMES)^hi^ PD1^hi^ subsets (17), but only the T-bet^hi^ PD1^mid^ subset, rather than EOMES^hi^ PD1^hi^, is reinvigorated by blockade of the PD1 pathway (18). In addition, the effectiveness of the PD1 pathway blockade depends on the pre-existing antitumor immune response in patients. In fact, targeting both PD1 and the CTLA4 pathway leads to adverse immune-related toxicities (19).

Therefore, here, virus- and tumor-specific exhausted CD8+ T cells were investigated to uncover the molecular mechanisms of CD8+ T cell dysfunction and also provide potential and highly efficient targets for therapeutic blockade in the treatment of cancer and chronic infection, as well as inflammation. Of particular note is that in mice models with cancer, antitumor immunity is enhanced via blocking multiple immune checkpoints(15, 20); thus, it is possible to make large achievements in immunotherapeutics if more targets are found. In this study, raw RNA-sequencing (RNA-seq) read datasets were collected from human and mouse samples, representing either normal or exhausted T cells. Potential immunosuppressive factors were revealed covering both widely reported and novel immune checkpoints, which may be considered prime immunotherapy targets.

## Materials and Methods

### Acquisition of the RNA-seq datasets

Raw RNA-seq reads of CD8+ T cells were collected from Gene Expression Omnibus (GEO). Three datasets were from *Homo sapiens* studies(21–23), and four datasets were from *Mus musculus*(24–27). Regarding the datasets, CD8+ T cell exhaustion occurs during non-small-cell lung cancer, hepatocellular carcinoma, type 1 diabetes (T1D), or chronic infection. Samples were further filtered using the following criteria: (i) CD8+ T cell samples were used; (ii) transgenic and PD1 blockade samples were excluded; (iii) in the T1D study, T cell immunoreceptor with Ig and ITIM domains (TIGIT)^+^ killer cell lectin like receptor G1 (KLRG1)^+^ [double high (DH)] and TIGIT^−^KLRG1^−^[double low (DL)] CD8+ T cells were used after 6 months of teplizumab antibody treatment, since the authors investigated that DH cells exhibited an exhaustion phenotype compared to DL cells. Moreover, DH cells treated with poliovirus receptor-Fc (a ligand for TIGIT) were excluded because they were downregulated. The accession numbers reported in this paper were GSE99531, GSE111389, GSE85530, GSE93006, GSE84820, GSE83978, and GSE86881.

### RNA-seq pipeline analysis

The raw RNA-seq reads were filtered using the FASTX-Toolkit fastq_quality_filter (28) by setting the parameters -q 20 (minimum quality score to keep) and -p 80 (minimum percent of bases that must have [-q] quality). Duplications caused by polymerase chain reaction amplification were removed such that paired- and single-end reads used fastuniq (28) and the clumpify.sh program from BBMap suite v38.32 (29), respectively. In particular, arbitrarily disordered files were repaired using the repair.sh program from BBMap. Pre-processed reads of humans and mice were then aligned to the GRCh38.94 (release 94) or GRCm38 (release 94) reference genome using STAR (30) and subsequently, expression was calculated using RNA-Seq by Expectation Maximization v1.3.1 (31). Transcripts per kilobase per million (TPM) values were used.

### Identification and analysis of differentially expressed genes (DEGs) between exhausted and non-exhausted CD8+ T cells

The TPM values in each sample were normalized using the up-quartile normalization method. The R package limma was used to remove batch effects. Principal component analysis (PCA) was performed before and after removing the batch effect via FactoMineR and factoextra packages in R. The RP function in the RankProd R package was used to analyze DEGs (32).

The topGene function in the package was used to identify DEGs that were required to pass a false discovery rate (FDR) threshold of 0.05, and a linear fold change threshold of 1.5. Gene ontology (GO) analysis based on the biological process and construction of protein-protein association networks was performed using the Search Tool for the Retrieval of Interacting Genes/Proteins (STRING, https://string-db.org/) database (33). A process or pathway was considered to be significantly enriched if the FDR value was < 0.05. The protein-protein interaction files obtained from STRING were then used to identify hub genes in the network. The hub genes with the top interaction degrees were extracted and analyzed using Cytoscape software version 3.7.2 (34) and its cytoHubba (35) plugin.

### Co-expression based on weighted correlation network analysis (WGCNA)

WGCNA (36) is an R package that is used to study clusters (modules) of highly correlated genes, the relationship between clinical traits and gene expression profiles, as well as calculating the correlation values within each module or module-trait relationships. The main WGCNA workflow was as follows: (i) The gene expression matrix and clinical traits were prepared and normalized TPM values were used. (ii) The soft thresholding power was determined by calling the network topology analysis pickSoftThreshold function. The set of candidate soft-thresholding powers ranged from 1–30. Then, the scale independence and mean connectivity for each power were calculated. A suitable power was selected if the degree of independence was more than 0.9 (human 11, mouse: 12). (iii) A co-expression network (unsigned network) was constructed and modules were identified. All genes were assigned to dozens of modules via hierarchical clustering and dynamic tree cutting. These modules were represented using unique color labels. (iv) Modules were related to external information, such as clinical traits. First, the module eigengene (ME) values were recalculated. An eigengene was the first principal component of a module expression matrix and thus was considered representative information for all genes in each module. Second, these eigengenes were correlated with clinical traits to obtain significant associations using the Pearson coefficient. Finally, the module-trait relationships were visualized using the labeledHeatmap function. (v) The weight was added to the existing module eigengenes and the relationships among eigengenes and the exhaustion trait were plotted using the plotEigengeneNetworks function, including a heatmap and dendrogram. (vi) The module membership (MM) was examined and the key drivers in interesting modules were identified. First, a matrix of Pearson correlation coefficients was obtained among genes and modules. Second, a similar matrix among genes and exhaustion traits was obtained. Third, the two matrices were combined and the interesting modules were analyzed and visualized using the verboseScatterplot function.

### Gene set enrichment analysis (GSEA)

GSEA was performed using the GSEA software version 4.0.2 (37). Eighty gene sets associated with T cell exhaustion traits were collected from the molecular signatures database v7 (MSigDB 7.0) and used for GSEA. Kyoto Encyclopedia of Genes and Genomes (KEGG) pathway enrichment analysis was performed using GSEA. The gene set used was c2.cp.kegg.v7.0.symbols.gmt. The normalized enrichment score and FDR values were calculated using permutation testing.

### Statistical analysis

Statistical analyses were performed using R software, version 3.5.2. Statistical analysis was performed using an unpaired two-tailed Student’s *t*-test (\**p* < 0.05, \*\**p* < 0.01, \*\*\**p* < 0.001).

## Results

### Human and mouse samples were characterized using RNA-seq datasets

The analysis procedure used in this study is shown in Figure 1. Raw RNA-seq reads were collected from seven studies, including three *Homo sapiens* and four *Mus musculus* studies. CD8+ T cell samples with exhausted or non-exhausted traits were selected according to rigid criteria (see the Materials and Methods). A total of 88 human and 35 mouse samples showing either exhausted or non-exhausted traits were used to comprehensively uncover the molecular mechanisms of CD8+ T cell dysfunction and identify highly efficient targets for the treatment of cancer and chronic infections (Table 1). Detailed information of the 123 samples is shown in Supplementary Table 1.

**Figure 1.**
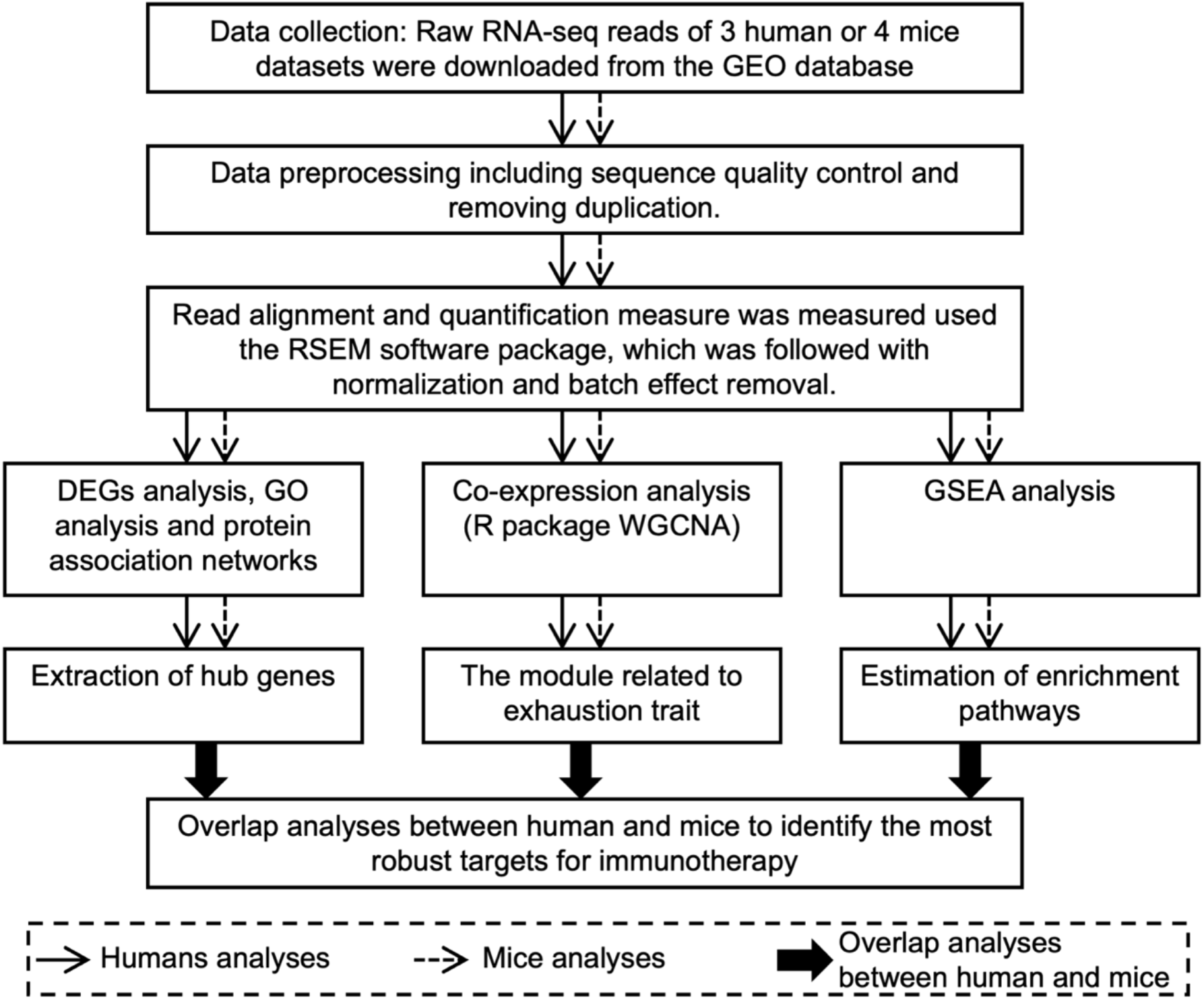
A schematic diagram of the workflow. GEO, Gene Expression Omnibus; RSEM, RNA-Seq by Expectation–Maximization; DEGs, Differentially expressed genes; GO, Gene ontology; WGCNA, weighted correlation network analysis; GSEA, gene set enrichment analysis.

**Table 1.** *Homo sapiens* and *Mus musculus* sample category information.

| Species | Status | Sample size | Reference |
| --- | --- | --- | --- |
| *Homo sapiens* | Exhausted | 11 | Thommen et al. (2018) (21) |
|  | Non-exhausted | 26 |  |
|  | Exhausted | 6 | Kim et al. (2018) (22) |
|  | Non-exhausted | 12 |  |
|  | Exhausted | 16 | Long et al. (2016) (23) |
|  | Non-exhausted | 17 |  |
| *Mus musculus* | Exhausted | 6 | Mognol et al. (2017) (24) |
|  | Non-exhausted | 12 |  |
|  | Exhausted | 2 | Man et al. (2017) (25) |
|  | Non-exhausted | 4 |  |
|  | Exhausted | 3 | Utzschneider et al. (2016) (26) |
|  | Non-exhausted | 3 |  |
|  | Exhausted | 5 | Pauken et al. (2016) (27) |
|  | Non-exhausted | 0 |  |

In this study, virus- and tumor-specific CD8+ T cells became exhausted within infections, cancer, or other disease microenvironments, such as LCMV infection, non-small-cell lung cancer, hepatocellular carcinoma, and T1D. Raw reads were then aligned to human or mouse reference genomes separately and TPM values were calculated (see the Materials and Methods section for more information). To compare gene expression profiles of exhausted and non-exhausted CD8+ T cells that were collected from different studies and diseases, batch effects were removed after TPM value normalization. PCA analysis was further performed to compare overall profiles before and after batch effect removal and normalization (Supplementary Figure 1 and 2). Original datasets with different resources were classified into different clusters for humans and mice; however, there were no obvious clusters after batch effect removal and normalization. Samples with different exhausted and non-exhausted traits were divided into different clusters after batch effect removal and normalization, indicating that the gene expression profiles showed differences due to these traits.

### DEGs of exhausted and non-exhausted CD8+ T cells

Having characterized the exhausted and non-exhausted CD8+ T cells, DEGs associated with exhausted CD8+ T cells were identified using normalized TPM values. Genes showing significantly different expression were defined as fold change > 1.5, and FDR < 0.05. A total of 199 upregulated and 441 downregulated genes were identified in exhausted CD8+ T cells in humans (Supplementary Table 2). Distinct expression profiles of these DEGs in both exhausted and non-exhausted CD8+ T cells were displayed and showed distinct patterns between exhausted and non-exhausted samples (Figure 2A). Compared to humans, 546 upregulated and 575 downregulated genes were identified in exhausted CD8+ T cells of mice (Supplementary Table 2). Subsequently, a heatmap of the 1,121 DEGs was constructed and showed obvious gene signatures in exhausted datasets, which were markedly distinct from non-exhausted datasets (Supplementary Figure 3A).

**Figure 2.**
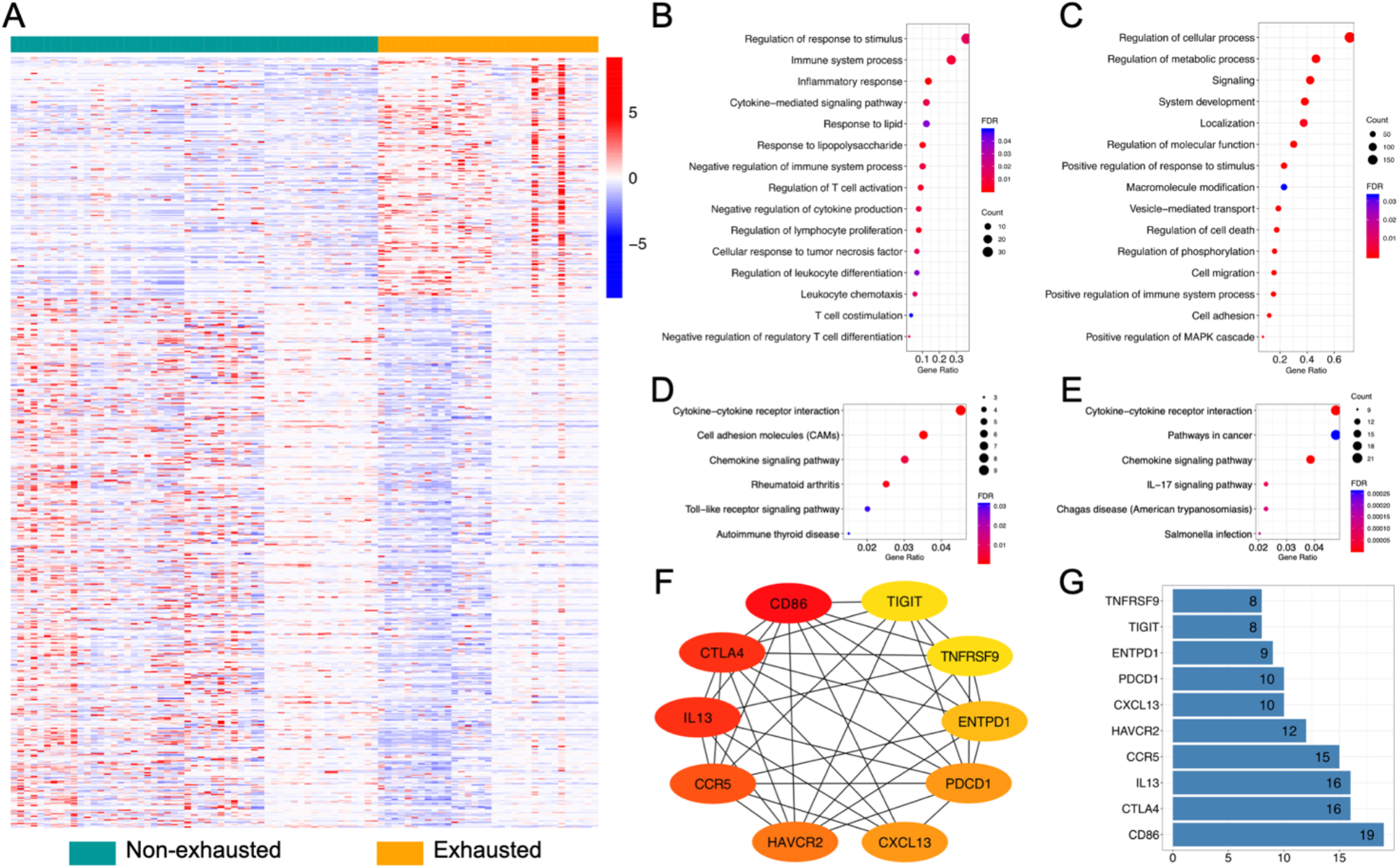
Human differentially expressed gene (DEG) analysis. (A) Heatmap constructed using 199 upregulated and 441 downregulated genes in exhausted T-cells. Significant changes are determined as fold change > 1.5 and false discovery rate (FDR) < 0.05. (B–C) Gene ontology (GO) and (D–E) Kyoto Encyclopedia of Genes and Genomes (KEGG) pathway enrichment analysis for human DEGs. Selected key biological processes in (B) upregulated and (C) downregulated genes. GO analysis is performed based on biological process. (D) All six enriched KEGG pathways in upregulated genes. (E) The top six enriched KEGG pathways in downregulated genes. GO terms and KEGG pathway remain when the FDR is < 0.05. (F–G) Functional protein association networks of upregulated genes. (F) Cytoscape software and Cytohubba plugin have been used to identify hub genes with top 10 interaction degrees via analyzing interactions files obtained from the Search Tool for the Retrieval of Interacting Genes/Proteins (STRING) database. (G) Interaction degrees of the hub genes. X-axis represents the number of adjacent genes.

Next, the DEGs in exhausted CD8+ T cells were used to identify enriched GO terms (biological processes) and KEGG pathways (Figure 2B–E, Supplementary Figure 3B–C). Upregulated genes were involved in several GO biological processes, such as regulation of cytokine production, lymphocyte proliferation, and regulatory T cell differentiation. This might be because, compared to normal tissues, T cells lose their proliferative potential and cytokine production capacity during exhaustion due to chronic infection or cancer microenvironments (38). Of particular note, the genes involved in the regulation of response to the stimulus, immune system process, and inflammatory response were among the most upregulated genes in exhausted T cells, including the inhibitory receptors PD1, CTLA4, HAVCR2 (also known as T-cell immunoglobulin, and mucin-domain containing-3 [TIM3]), and TIGIT. Downregulated genes were enriched in cell migration, cell adhesion, and regulation of phosphorylation. In particular, exhausted CD8+ T cells display suppressed IFN-γ production(2, 39), whereas here, its receptor, IFN-γ receptor 2 (IFNGR2), is expressed at a low level among exhausted samples; thus, we hypothesized that this finding might further enhance the T cell exhaustion trait. Functional enriched KEGG pathway analysis indicates that human DEGs are enriched in several pathways, as shown in Figure 2D–E. Among the total enriched pathways of upregulated genes, cytokine-cytokine receptor interaction, rheumatoid arthritis, and chemokine signaling pathways were also enriched in downregulated genes. Cell adhesion molecules, autoimmune thyroid disease, and Toll-like receptor signaling pathways were unique to upregulated genes. Intriguingly, the IL-17 signaling pathway is predominantly enriched in downregulated genes, which is rarely explained in exhausted T cells, as the production of various interleukins is lost in these cells, such as IL-2, IL-7, and IL-15 (2).

Additionally, as immune checkpoint inhibitors were overexpressed in exhausted T cells, association networks of upregulated genes were constructed using the STRING website (Supplementary Figure 4A and 5A). As expected, most edges in the network were derived from co-expression (shown as yellow-green), while text-mining information (shown in black) also supported a significant portion of the edges, which were associated with two main networks for both humans and mice. One network was related to the immune system process, containing genes, such as PD1, TIGIT, HAVCR2, and TNF superfamily member 4 (TNFSF4). Another main network, which contained interactions supported by experiments (shown in magenta), was associated with the mitotic cell cycle process and cell population proliferation, such as maternal embryonic leucine zipper kinase (MELK) and mitotic checkpoint serine/threonine-protein kinase BUB1 beta (BUB1B), encoded by *MELK* and *BUB1B*, respectively. Proteins that belonged to neither of the two main networks (possibly because of the lower number of studies compared to PD1) and associated with the immune system process, such as adhesion G protein-coupled receptor G1 (ADGRG1), thymocyte selection associated high mobility group box (TOX/TOX2), and CD200 receptor 1 (CD200R/CD200R1), might be used to identify novel potential targets for immunotherapy. To identify the key drivers of the main network associated with the immune system process, the Cytoscape Cytohubba plugin was used to identify hub genes with the top 10 interaction degrees by analyzing interactions files obtained from the STRING database (Figure 2F–G, Supplementary Figure 3D–E). Among the top 10 hub genes in humans and mice, three of the key driver genes, programmed cell death protein 1 (*PDCD1*), *HAVCR2*, and *TNFRSF9* (its alternative name is CD137), were simultaneously identified in humans and mice.

### Identification of key genes in exhausted CD8+ T cells based on WGCNA

WGCNA is an effective method to detect co-expressed modules and hub genes in many aspects (40). To investigate the relationship between the modules and clinical traits and identify potential prognostic markers of exhausted T cells, co-expressed modules were constructed with WGCNA using the gene expression profiles of exhausted and non-exhausted CD8+ T cells in humans (Figure 3) and mice (Supplementary Figure 6). A total of 11,499 genes from 88 human samples were used for WGCNA. A suitable power of 11 was selected for the co-expression analysis. Genes with common biological functions and associations were assigned into 24 co-expressed modules (clusters), including the gray module, which contained the genes that were not classified into any of the other 23 modules. Subsequently, the correlation values between the modules and clinical traits (exhausted or non-exhausted) were calculated based on the correlations between MEs and clinical traits, demonstrating that the red module was most positively correlated with exhaustion and that the correlation value was 0.52 (mouse: turquoise module, 0.71). To detect groups of correlated eigengenes and exhaustion, the existing 23 modules (except the gray module) were further analyzed using eigengene dendrogram and heatmap. The dendrogram also suggested that the red module was highly related to T cell exhaustion. Taken together, the genes in the red modules, including 263 genes, might play a critical role in T cell exhaustion. Next, to measure the importance of each gene within a specific module and their contributions to the phenotype, the MM vs. gene significance (GS) of the interesting red module was calculated. Notably, the well-studied inhibitory receptors, PD1, CTLA4, HAVCR2, TIGIT, and CD27 (a member of the tumor necrosis factor receptor superfamily), showed a high level of MM vs. GS, indicating that they were highly correlated with T cell exhaustion and key components of the underlying biological function.

**Figure 3.**
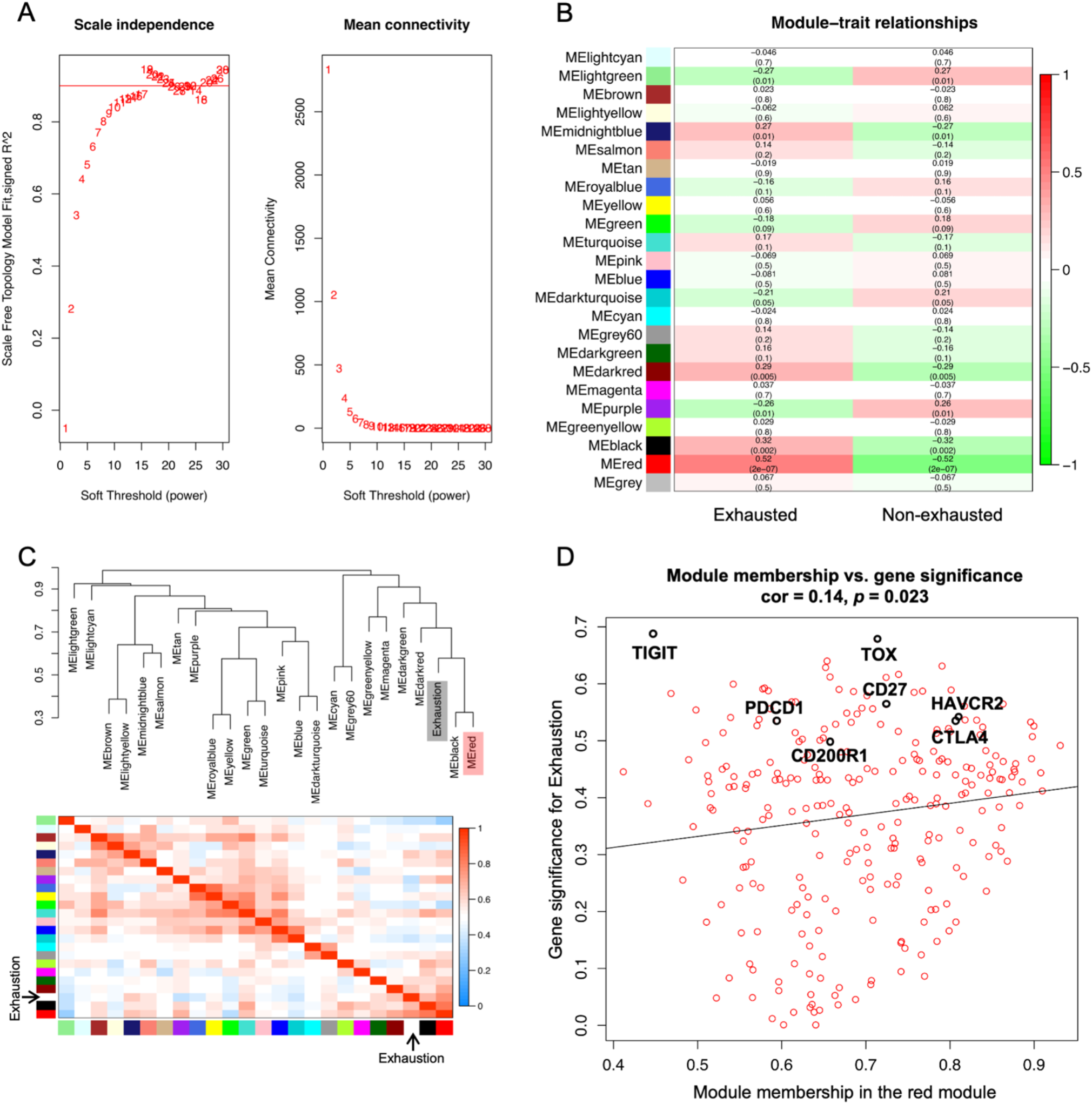
Co-expression analysis in human exhausted T-cells. (A) Analysis of a set of soft thresholding powers. The left panel is the scale free fit index as a function of soft thresholding power. The right panel is mean connectivity as a function of the soft thresholding power. (B) Heatmap of module and trait correlation. Module is marked by unique color labels. ME: module eigengene. (C) Eigengene dendrogram and heatmap between modules and the exhaustion trait. (D) Scatterplot of gene significance for exhaustion trait (y-axis) vs. membership in a selected module (x-axis).

Genes in the red module were further used for GO (biological process) and KEGG pathway enrichment analyses (Figure 4). Metabolic derangements occur in exhausted T cells (2). Here, the GO and KEGG pathway analyses demonstrated that these genes were particularly enriched in metabolic pathways (such as pyrimidine metabolism and glycine, serine, and threonine metabolism), cell cycle, DNA replication, phosphorylation, and p53 signaling pathway, which may shed light on T cell exhaustion mechanisms.

**Figure 4.**
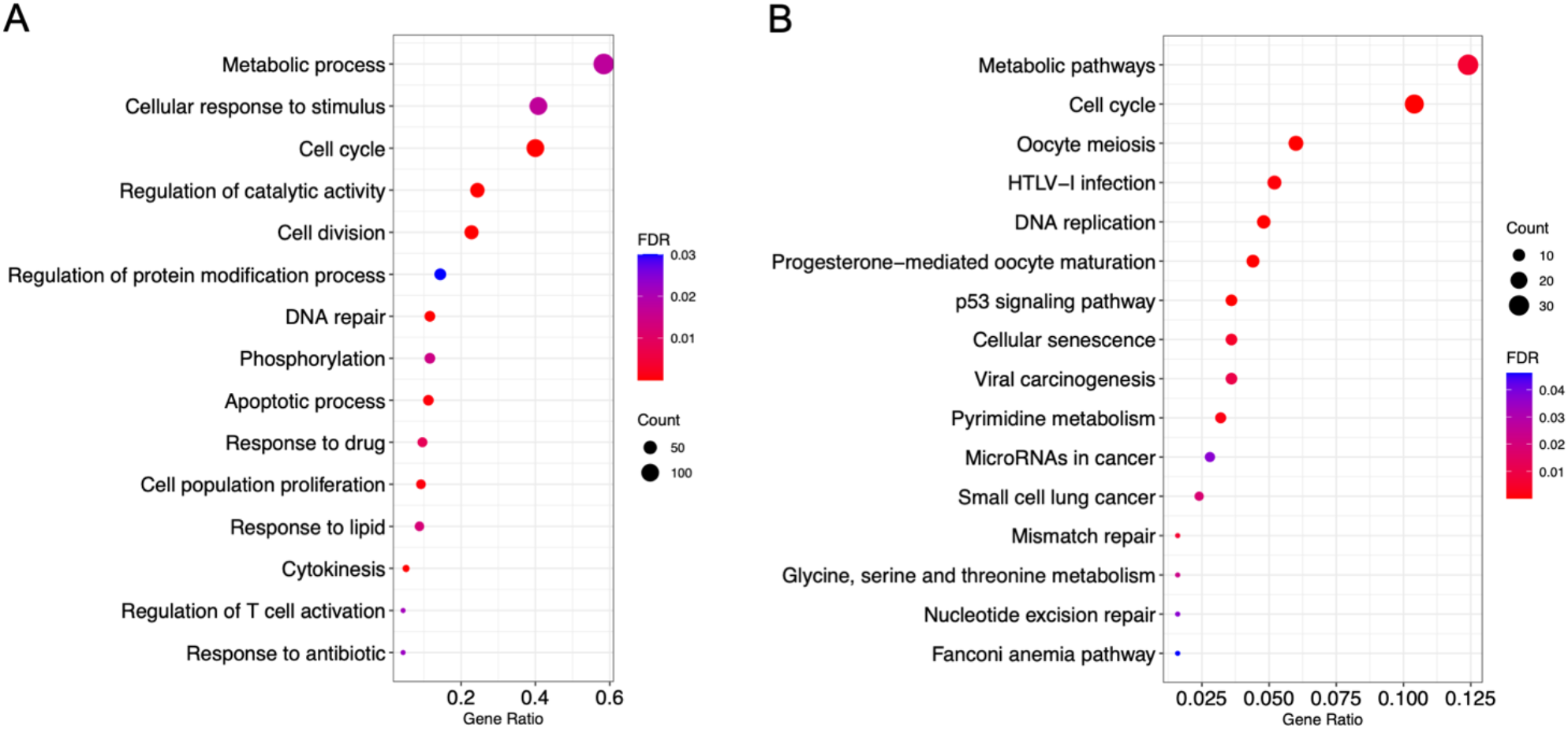
Gene ontology (GO) and Kyoto Encyclopedia of Genes and Genomes (KEGG) pathway enrichment analysis for human red module genes. (A) Selected key biological processes. (B) All enriched KEGG pathways. GO terms and KEGG pathway remain when the FDR is < 0.05.

### TOX and CD200R1 as potential targets

Since antibodies targeting immune checkpoint inhibitors are usually overexpressed in exhausted T cells, reversing T cell exhaustion is a promising approach for cancer immunotherapy. Thus, the above findings from the upregulated genes and co-expression analyses (Figure 5A) were integrated to discover potential and robust therapeutic targets. First, 21 overlapping upregulated genes were identified between humans and mice. Second, 175 overlapping genes were identified from the most exhausted-correlated modules (human, red module; mouse, turquoise module) based on co-expression analysis. Eventually, nine overlapping genes were obtained by combining the upregulated genes and co-expression analyses between humans and mice. The nine overlapping genes were further used to construct protein-protein association networks using the STRING database. Strikingly, six of the nine proteins [PD1 (encoded by *PDCD1*), TIGIT, HAVCR2, TNFSF4 (also known as CD134 or OX40L), TOX, and CD200R1] were associated with the regulation of immune system processes. In particular, the well-studied PD1, TIGIT, HAVCR2, and TNFSF4 were classified into one of the main networks, while TOX and CD200R1 belonged to neither of the main networks and require further study. Of particular note, *TOX* and *CD200R1* showed a high level of MM vs. GS, suggesting that these two genes were most important during T cell exhaustion (Figure 3D). In addition, the expression profiles of the six genes were confirmed at the mRNA level (Figure 5B–C). Compared to non-exhausted CD8+ T cells, the expression of *TOX* and *CD200R1* was similar to that of the well-studied inhibitory receptors, which showed distinctly high levels of expression in exhausted CD8+ T cells. Taken together, the meta-analysis of human and mouse exhausted CD8+ T cells suggested that TOX and CD200R1 may be prime and robust targets in the treatment of cancer and other inflammatory settings.

**Figure 5.**
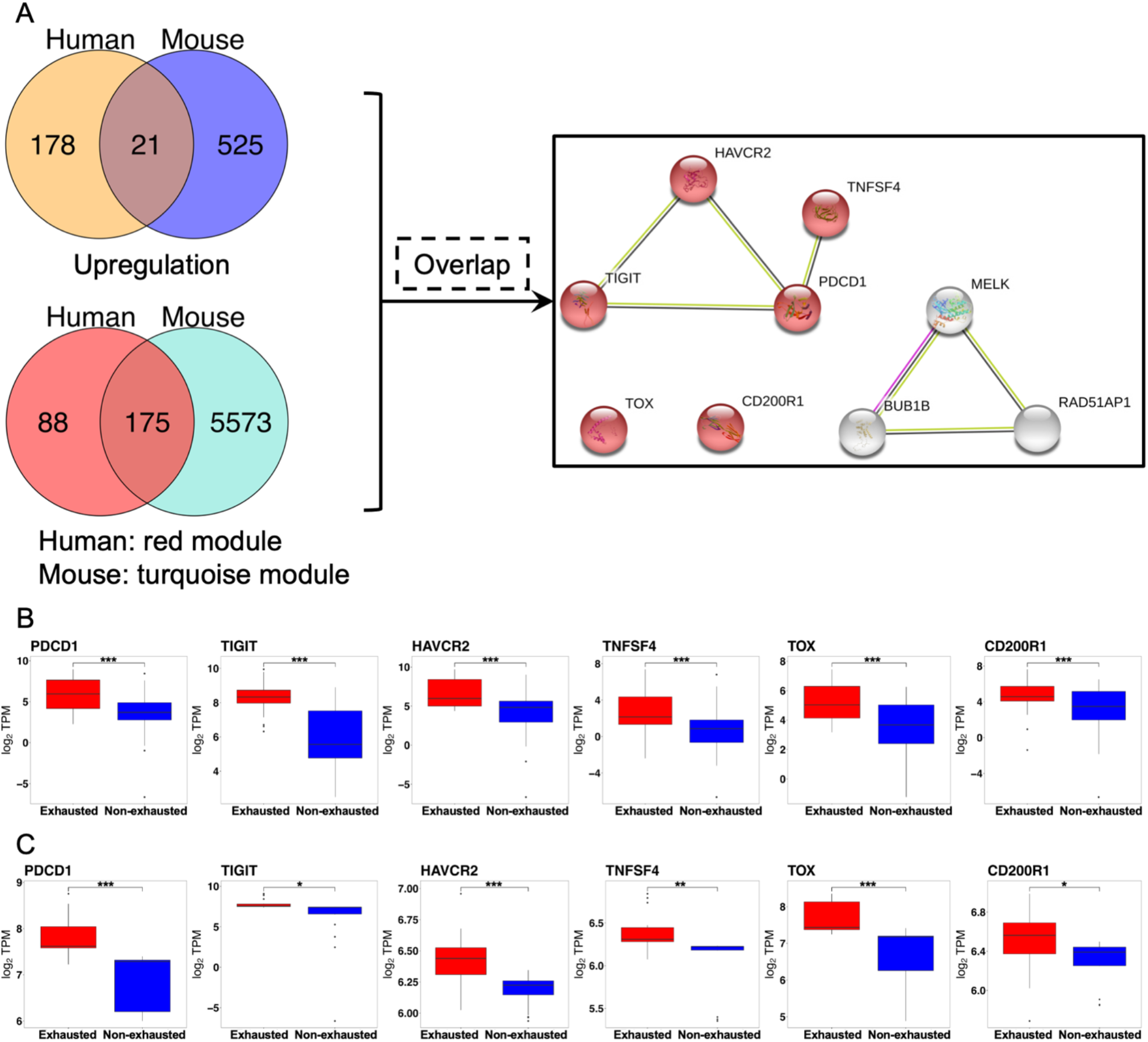
Overlap genes between human and mouse sets. (A) Overlap of upregulated genes in exhausted T-cells and most correlative modules with exhaustion between human and mouse, as well as protein associated network of nine proteins in both the overlap of upregulated genes and selected modules. Red colored nodes: regulation of immune system process (GO:0002682). (B) and (C) show the expression of selected genes at the mRNA level in human and mouse datasets, respectively. Median values are indicated by lines in the box and whisker plot. Hinge values and whisker 1.5* interquartile range (IQR) values have been calculated using Tukey. Statistical analysis has been performed using an unpaired two-tailed Student’s *t*-test (\**p* < 0.05, \*\**p* < 0.01, \*\*\**p* < 0.001).

### Predefined exhaustion gene sets were enriched in exhausted CD8+ T cells

Additionally, to interpret genome-wide transcriptional profiles of the exhausted T cell samples in this study and further examine the molecular mechanisms of T cell exhaustion, GSEA was performed using the 80 predefined gene sets collected from MSigDB that contained information on immunologic signatures, and gene sets from the biomedical literature. The 80 gene sets were associated with CD8+ T cell exhaustion during chronic infection or cancer. Compared to 17 gene sets in mice, 12 of 80 gene sets were significantly enriched in exhausted CD8+ T cells in humans (*p* < 0.05) (Supplementary Table 3). The gene members of the majority of those enriched gene sets were either upregulated or downregulated in comparison to exhausted or PD1 high CD8+ T cells versus effector, naïve, PD1 low, or memory CD8+ T cells. Four enriched gene sets in the exhausted CD8+ T cells were shared between humans and mice (Figure 6 and Supplementary Figure 7), including gene sets GSE9650_effector_VS_exhausted_CD8_TCELL_DN (genes downregulated in comparison to effector CD8 T cells versus exhausted CD8 T cells), GSE9650_naïve_VS_exhausted_CD8_TCELL_DN (genes downregulated in comparison to naive CD8 T cells versus exhausted CD8 T cells), GSE9650_effector_VS_memory_CD8_TCELL_UP (genes upregulated in comparison to effector CD8 T cells versus memory CD8 T cells), GSE9650_exhausted_VS_memory_CD8_TCELL_UP (genes upregulated in comparison to exhausted CD8 T cells versus memory CD8 T cells). These gene sets were obtained from the MSigDB C7 immunologic signatures (41). Most of the genes in the four gene sets were categorized as transcription factors, cell differentiation markers, oncogenes, cytokines, and growth factors. Additionally, many DEGs of human exhausted CD8+ T cells were identified in the enriched gene sets, such as *PDCD1* and *CTLA4*. In summary, these findings partially demonstrated the mechanisms of T cell exhaustion following chronic infection or cancer, especially the loss of cytokine secretion. The common signature between humans and mice may provide a framework for therapeutic interventions targeting exhausted CD8+ T cells.

**Figure 6.**
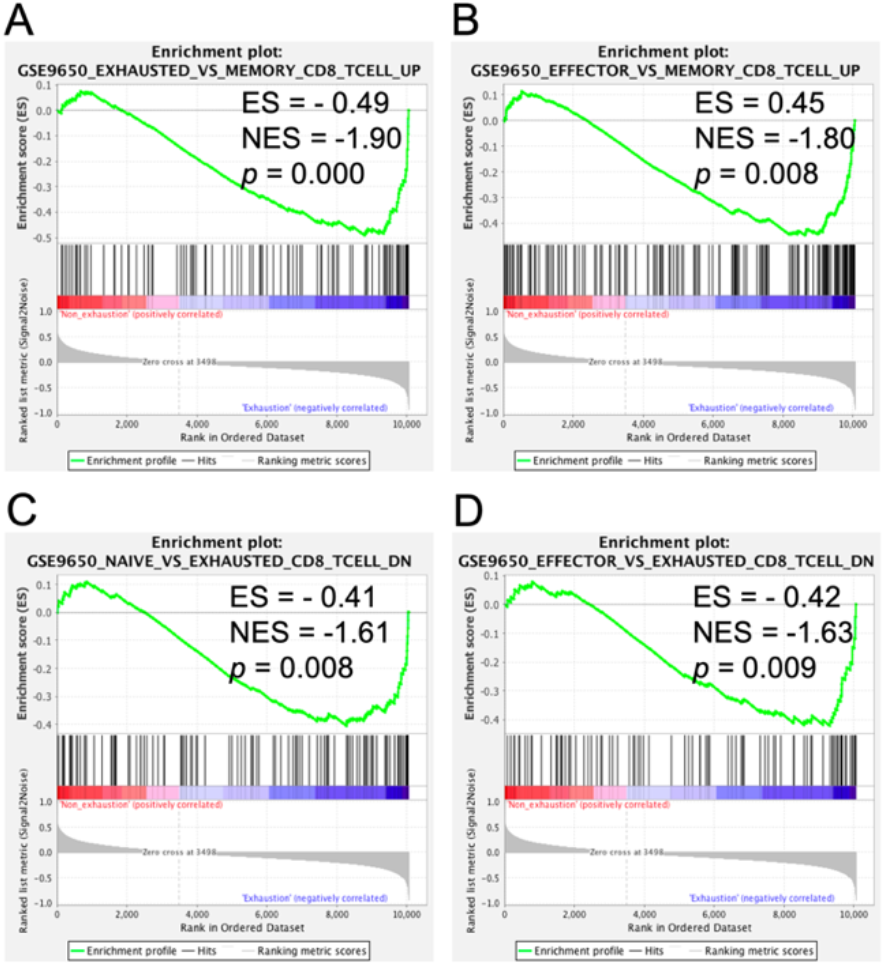
Gene set enrichment analysis (GSEA) of exhausted versus non-exhausted CD8+ T-cells. (A–D) The four shared enriched gene sets in human exhausted T-cells with a *p*-value < 0.05. ES, enrichment score; NES, normalized enrichment score.

## Discussion

Immunotherapies that reactivate exhausted T cells by targeting immune checkpoint inhibitors have emerged as promising treatment options (15, 16, 42, 43). Understanding the features of and pathways to T cell exhaustion has crucial implications for successfully blocking immune checkpoint inhibitors and immunotherapies. However, therapeutic efficacy remains low, and T cell exhaustion mechanisms are yet to be fully described. As T cell exhaustion has been separately described in humans and mice during chronic infection, cancer, HIV, or hepatitis C virus infection (9–11, 38), in this study, both human and mouse exhausted CD8+ T cells were comprehensively analyzed. As exhaustion may display potential differences between cancer and chronic infection, the exhausted T cells used in this study were developed during multiple microenvironments, such as non-small-cell lung cancer, hepatocellular carcinoma, T1D, or chronic infections with seven origins. Above all, these data might be able to identify more robust targets for therapies and shed light on the mechanisms of exhaustion.

The independent meta-analyses of humans and mice indicated that two inhibitory receptors, TOX and CD200R1, were significantly upregulated in exhausted CD8+ T cells and were key drivers of the modules that contributed to the exhaustion trait based on the WGCNA. Thus, TOX and CD200R1 are prime and robust targets for the treatment of multiple cancers and chronic infections in both mice and humans. Recent studies have also highlighted the important role of the TOX transcriptional regulator in driving T cell exhaustion and diminishing anti-tumor or anti-viral functions(44–46). CD200R1 is also considered a viable therapeutic target in cancer by blocking CD200R in the CD200R1 pathway (43). Compared to TOX and CD200R1, G protein-coupled receptor 56 (GPR56) might also be a candidate target, even though its robustness is weaker. GPR56 was upregulated in both human and mouse exhausted T cells, but was only co-expressed in the selected mouse turquoise module, and not in the human red module. GPR56 is encoded by *ADGRG1* (47) and is expressed in all human cytotoxic lymphocytes (48). When blocking inhibitor receptors as a immunotherapeutic, antibody targeting of multiple inhibitory receptors enhances antitumor immunity and expands efficacy (15). Therefore, the potential targets identified in this study may provide more opportunities for immunotherapy when using combinatorial approaches.

Additionally, the mechanisms underlying T cell exhaustion were uncovered. Hallmarks of exhausted T cells include overexpression of inhibitory receptors, loss of proliferative potential and cytokine production, markedly different transcriptional profiles, metabolic derangement, and the use of key transcription factors (2, 11, 38, 41, 49, 50). These findings might explain and demonstrate the formation of these key characteristics. First, the inhibitory receptors, PD1, CTLA4, TIGIT, HAVCR2, and TNFSF4 were upregulated. Second, several biological pathways were modulated via either significantly upregulated or downregulated genes, which potentially indicated impaired T cell capacity to proliferate and secrete cytokines. Specifically, the GO and KEGG pathway enrichment analyses suggested that DEGs were enriched in cytokine-cytokine receptor interaction, immune system process, cell cycle, phosphorylation, and the IL-17 signaling pathway. Of note, only one pathway, cytokine-cytokine receptor interaction, was enriched in mouse upregulated genes. In addition, the enriched IL-17 signaling pathway in human downregulated genes might provide a novel way to understand T cell exhaustion. Third, co-expressed genes based on WGCNA demonstrated altered metabolism in exhausted T cells. Only protein-protein interaction network and KEGG enrichment analyses of selected human red modules were performed, as there were too many genes in the mouse turquoise module (5748 genes). Thus, genes in the human red module were enriched in metabolic pathways, such as pyrimidine metabolism and glycine, serine, and threonine metabolism. In fact, cell cycle, DNA replication, nucleotide excision repair, and p53 signaling pathways were enriched in genes in the red module. Moreover, genes associated with cell cycle regulation and mitotic checkpoint, including cyclin dependent kinase 1, cyclin B1, budding uninhibited by benzimidazoles 1 (*BUB1*), cyclin A2, mitotic arrest deficient 2 like 1, *BUB1B*, cell division cycle 20, and cyclin B2 were the hub genes of the protein interaction network. Among the eight hub genes, only *BUB1* and *BUB1B* were significantly upregulated.

Additionally, according to the GSEA, a minority of the upregulated or downregulated genes identified in this study were observed in the gene members of the four shared gene sets. Specifically, to our surprise, upregulated *ADGRG1* was identified from two shared gene sets, effector_VS_exhausted_CD8_TCELL_DN and exhausted_VS_memory_CD8_TCELL_UP, which supported our previous hypothesis that ADGRG1 was a potential target for treatment. The downregulated genes, annexin A1 (*ANXA1*) and interleukin 12 receptor, beta 2 subunit (*IL12RB2*) were observed in the gene set effector_VS_memory_CD8_TCELL_UP. ANXA1 and IL12RB2 play important roles in Th1 cell differentiation and are thought to contribute to inflammatory responses and host defense(51, 52). The legumain gene is downregulated in exhausted_VS_memory_CD8_TCELL_UP and may be involved in endogenous proteins for major histocompatibility class II presentation in the lysosomal/endosomal system(53–55).

Taken together, by taking advantage of the meta-analyses between humans and mice, a novel way to understand the mechanisms of T cell exhaustion was demonstrated and provided robust potential targets for the treatment of cancer and chronic infection. Although T cell exhaustion has been identified in humans and mice, it may show differences to some degree, such as altered transcriptional profiles and transcriptional factors. Thus, to obtain inhibitory receptor blockers in cancer immunotherapy using mice as models, further comprehensive meta-analyses of multiple datasets that include humans and mice will be required in the future.

## Acknowledgements

L. Z. and H. N. were supported by the Development of Key Technologies for Next-Generation Artificial Intelligence/Robots from the New Energy and Industrial Technology Development Organization, Japan.

## Notes

### Competing Interest Statement

The authors have declared no competing interest.

