## Supplemental Figures for "Meta-analysis of exhausted CD8+ T cells from *Homo sapiens* and *Mus musculus* provides robust targets for immunotherapy"

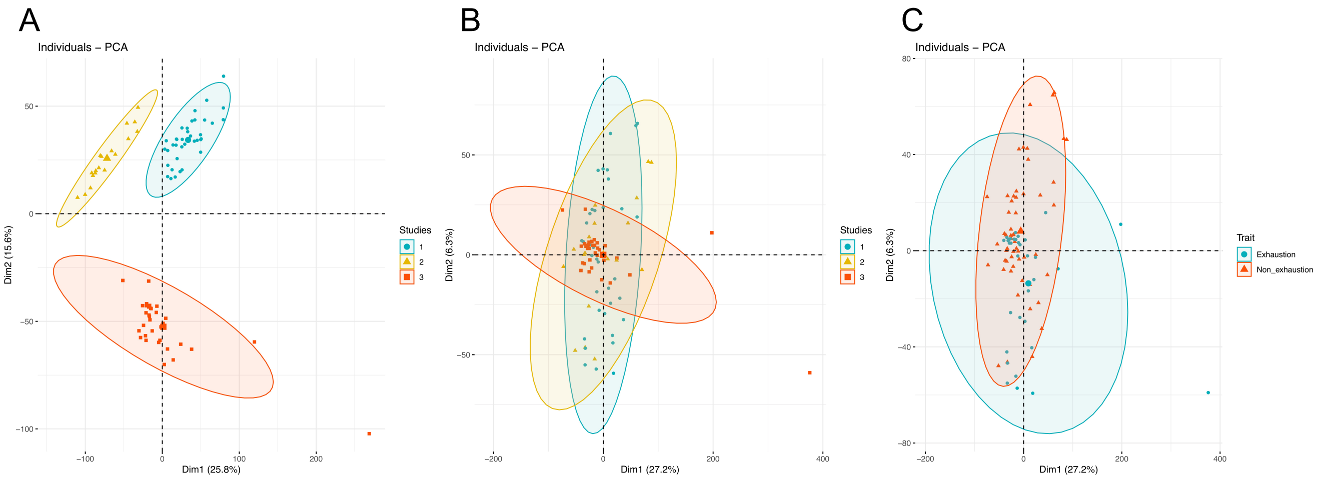


Supplementary Figure 1. Principal component analysis (PCA) of three human datasets.
(A) and (B) display clustering based on the dataset source before and after batch effect removal and normalization, respectively. (C) shows the clustering of non-exhausted and exhausted traits after batch effect removal.

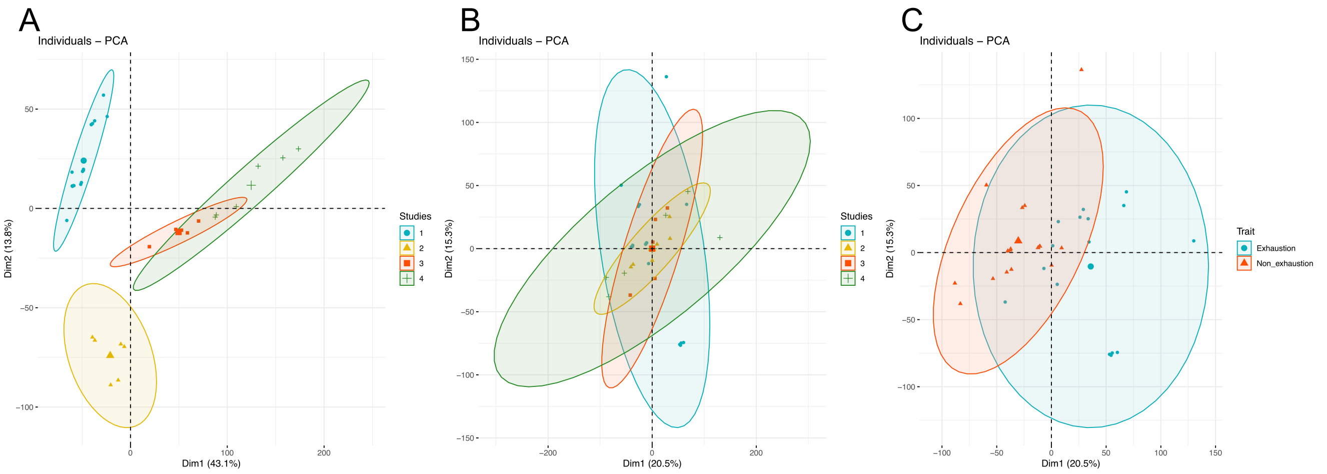


Supplementary Figure 2. Principal component analysis (PCA) of four mouse datasets.
(A) and (B) display clustering based on the dataset source before and after batch effect removal, respectively. (B) shows the clustering of non-exhausted and exhausted traits after batch effect removal.

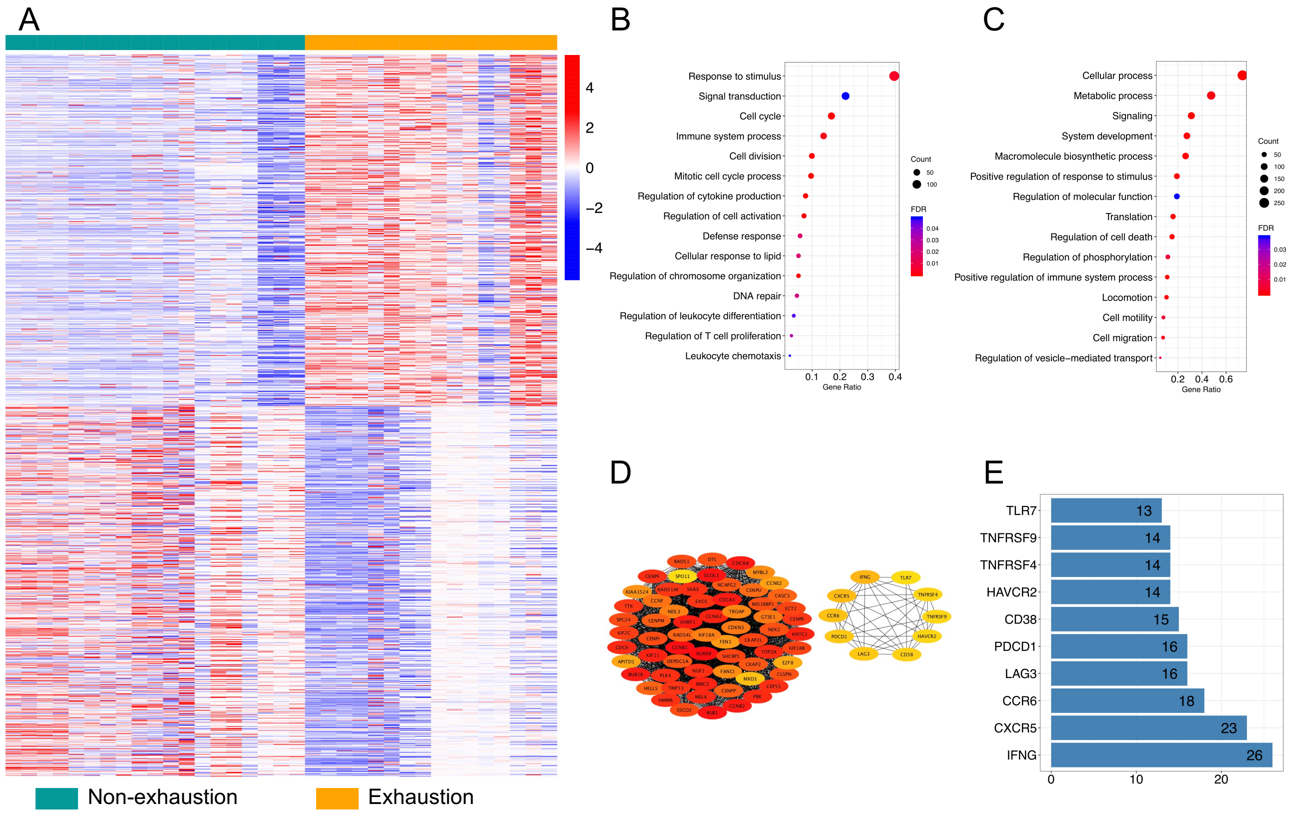


Supplementary Figure 3. Mouse gene expression profile of differentially expressed genes (DEGs).
(A) Heatmap constructed using 546 upregulated and 575 downregulated genes in exhausted T-cells. (B–C) Gene ontology (GO) enrichment analysis for mouse DEGs. Selected key biological processes in (B) upregulated and (C) downregulated genes. (D–E) Functional protein association networks of upregulated genes. Top 10 interaction degrees of hub genes are  displayed.

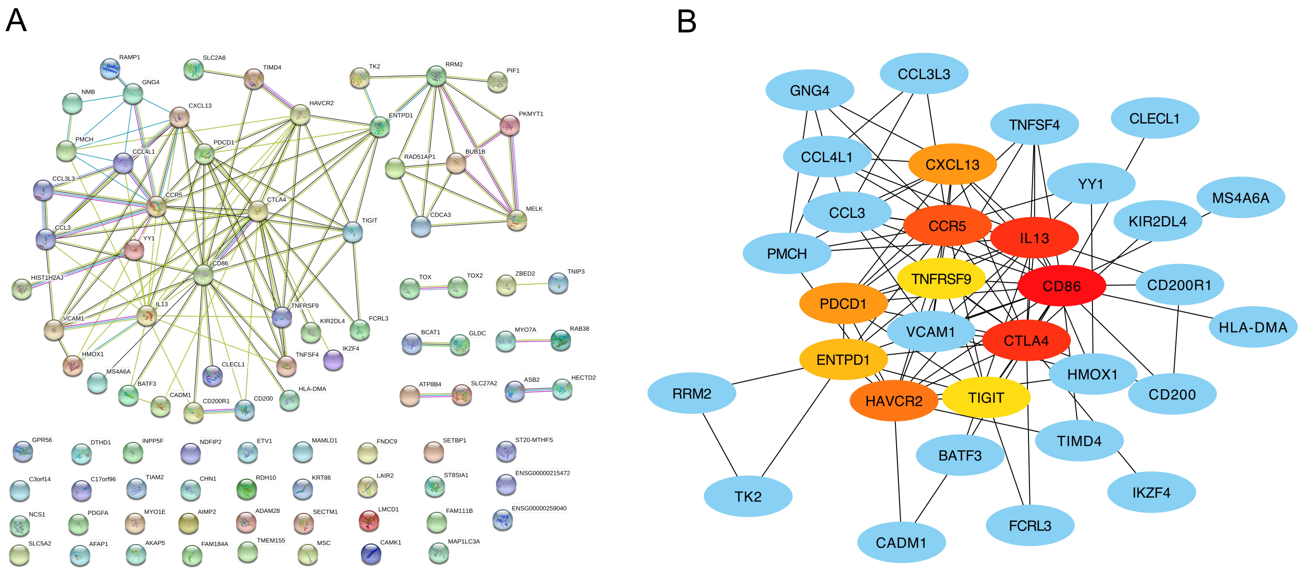


Supplementary Figure 4. Functional protein association networks of human upregulated genes.
(A) The network of upregulated genes constructed using the Search Tool for the Retrieval of Interacting Genes/Proteins (STRING) database. (B) Cytoscape software and Cytohubba plugin have been used to identify hub genes with top 10 interaction degrees via analyzing interactions files obtained from STRING.

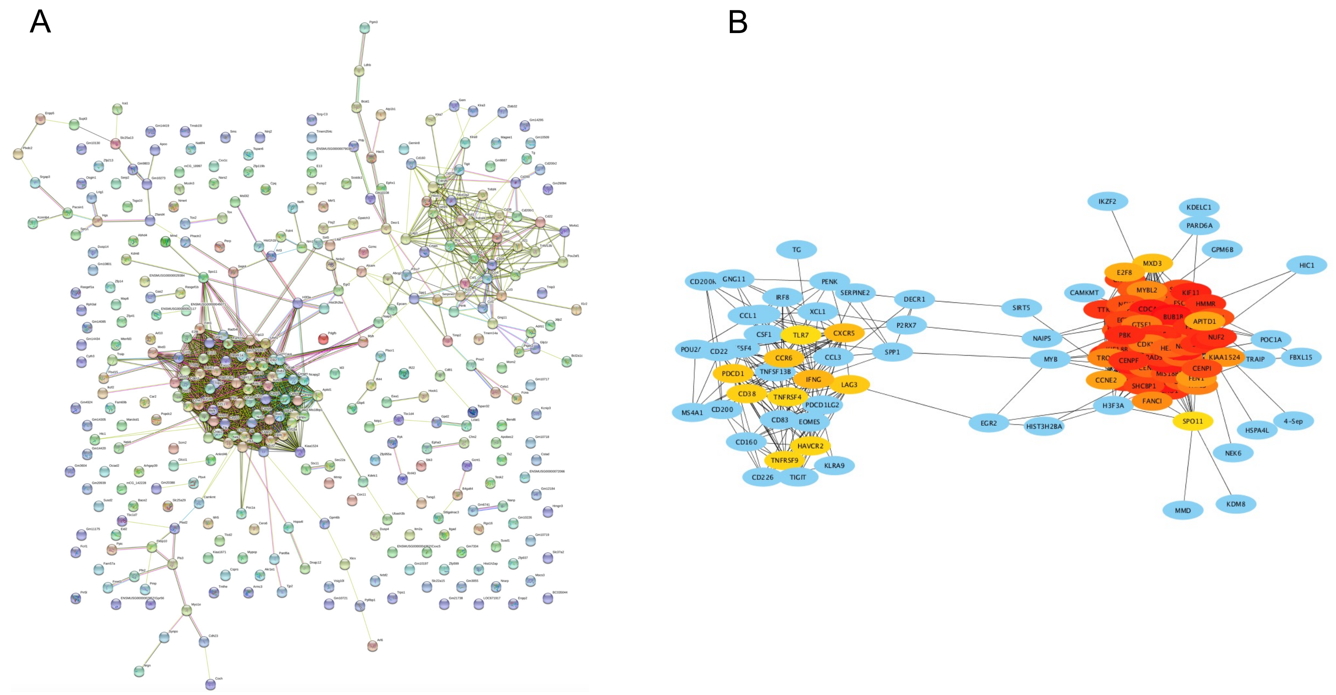


Supplementary Figure 5. Functional protein association networks of mouse upregulated genes.

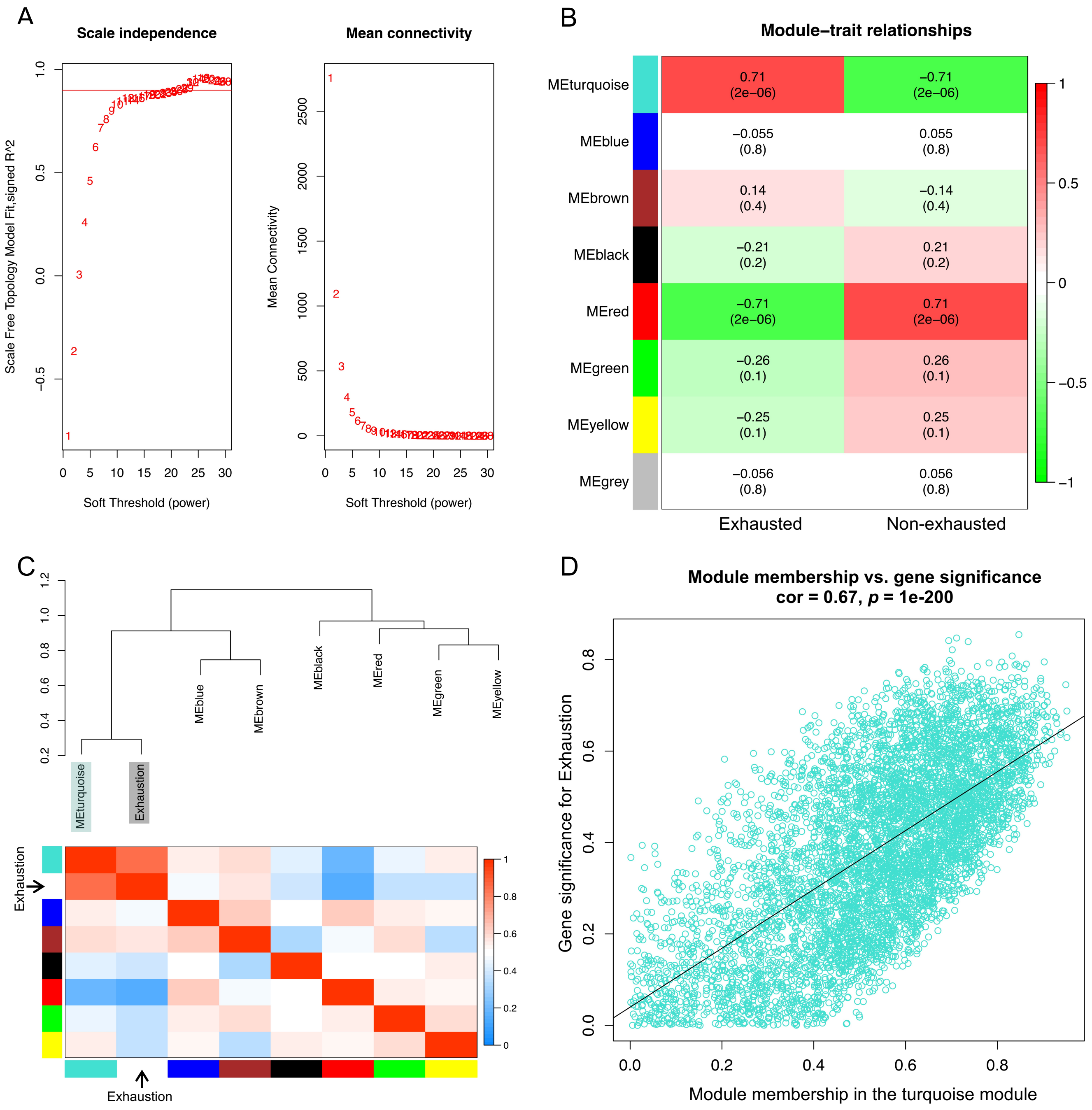


Supplementary Figure 6. Co-expression analysis of mouse exhausted T-cells.

(A) Analysis of a set of soft thresholding powers. (B) Heatmap of module and trait correlation. (C) Eigengene dendrogram and heatmap between modules and the exhaustion trait. (D) Scatterplot of gene significance for exhaustion trait (y-axis) vs. membership in a selected module (x-axis).

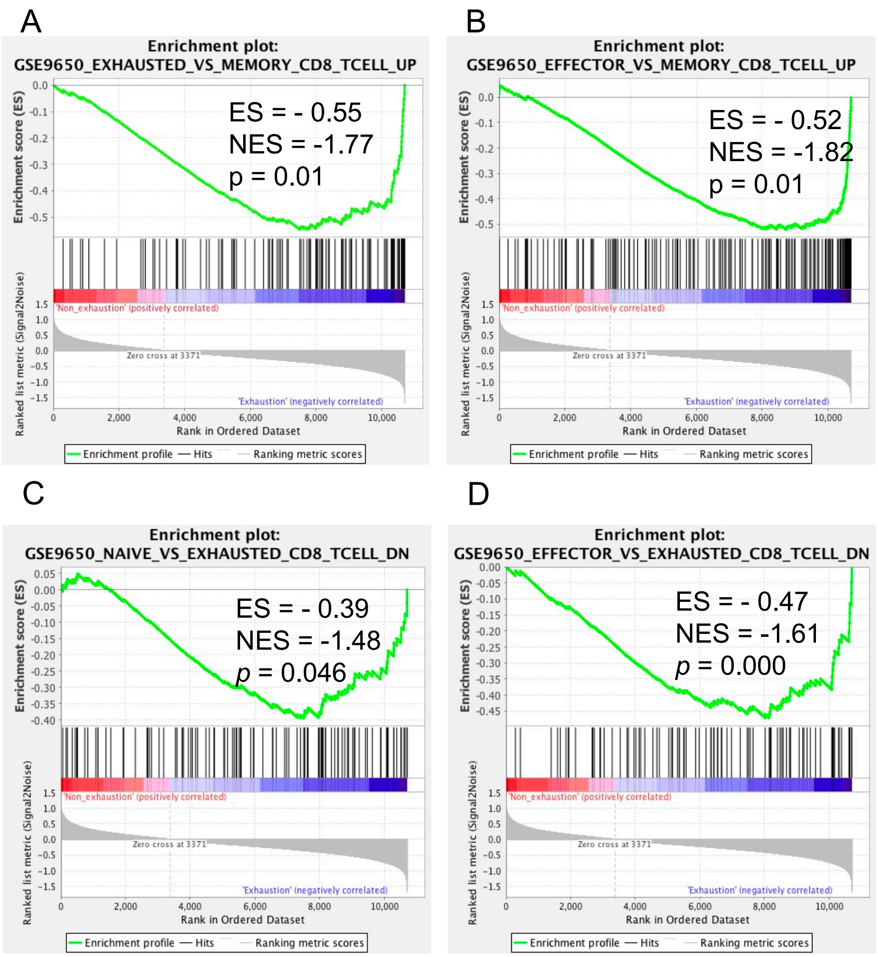


Supplementary Figure 7. Gene set enrichment analysis (GSEA) of exhausted versus non-exhausted CD8+ T-cells.
(A–D) The four shared enriched gene sets in mouse exhausted T-cells with a p-value < 0.05. ES, enrichment score; NES, normalized enrichment score.
